## Supplementary material for "UNC-6/Netrin promotes both adhesion and directed growth within a single axon": Document S1

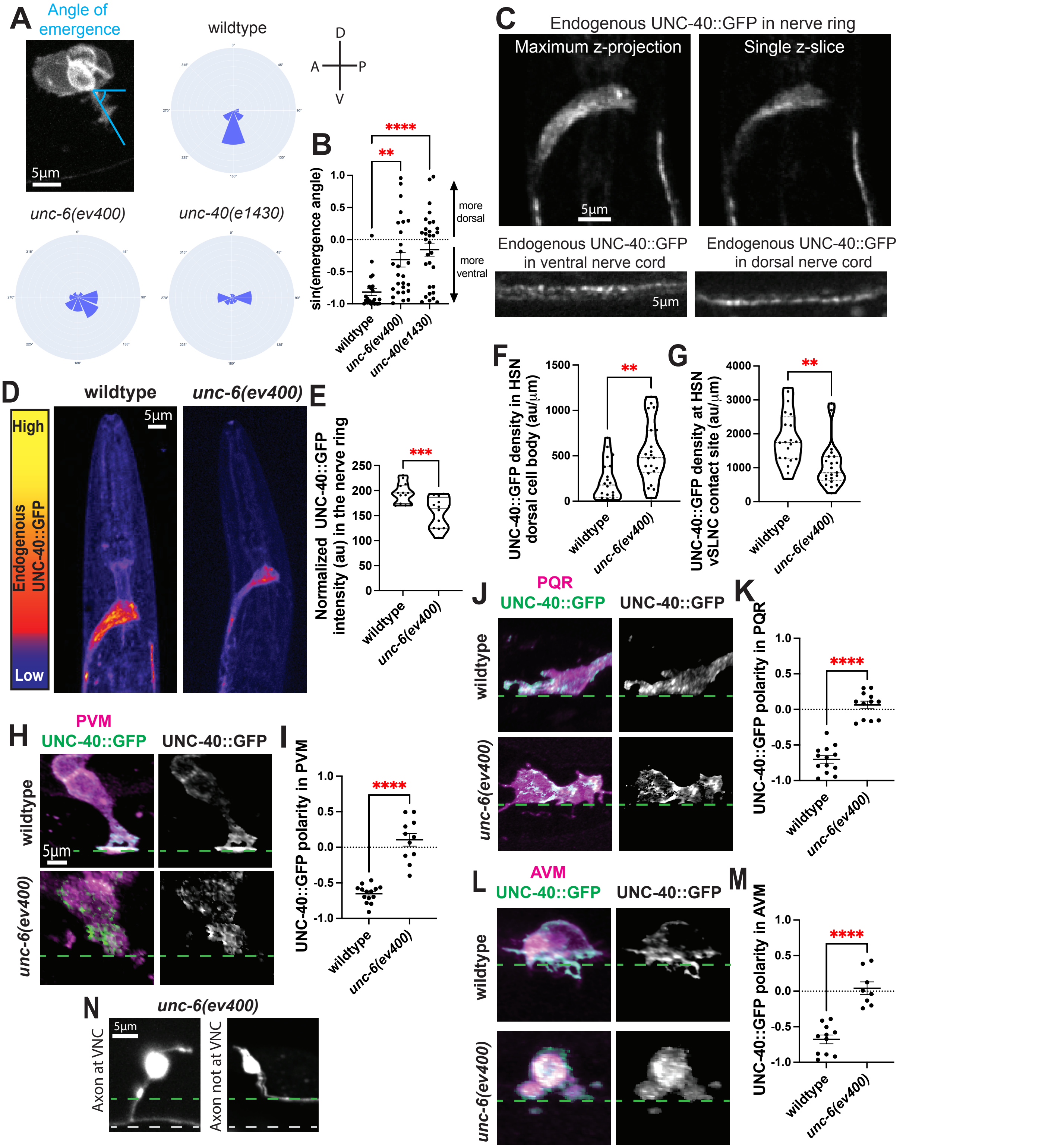


**Figure S1 Axon emergence and UNC-40 polarization are dependent on UNC-6, related to Figure 2.** A. Left: Confocal image of PDE axon emergence. Blue angle denotes measured angle of emergence relative to the animal's posterior. Right: Rose plots of PDE axon emergence. B. Graph comparing the sine function of axon emergence angle in wildtype, *unc-6(ev400)*, and *unc-40(e1430)*animals. 1 indicates 90° dorsal axon emergence and -1 indicates 90° ventral axon emergence. C. Confocal images of UNC-40::GFP in the nerve ring, VNC, and dorsal nerve cord. D. Confocal images of UNC-40::GFP in the nerve ring. E. Graph of fluorescent intensity of UNC-40::GFP in the nerve ring. E-F. Integrated UNC-40::GFP density along HSN dorsal cell body (E) and HSN contact site with vSLNC (F) in wildtype and *unc-6(ev400)* animals. H,J,L. Airyscan images of UNC-40::GFP in PVM (H), PQR (J), and AVM (L) after vSLNC contact. I,K,M. Graphs of UNC-40::GFP polarity PVM (I), PQR (K), and AVM (M). N. Confocal images of PDE adult morphology in *unc-6(ev400)* mutants . Left image shows an axon contacting the VNC and right image shows an axon that has not reached the VNC. Dashed green line denotes vSLNC and dashed gray line denotes VNC. SEM is shown. Scale bar is 5 µm in A,C,D,H,J,L,N.Ordinary one-way ANOVA with multiple comparisons is used in B. Unpaired t-test is used in E-G,I,K,M.

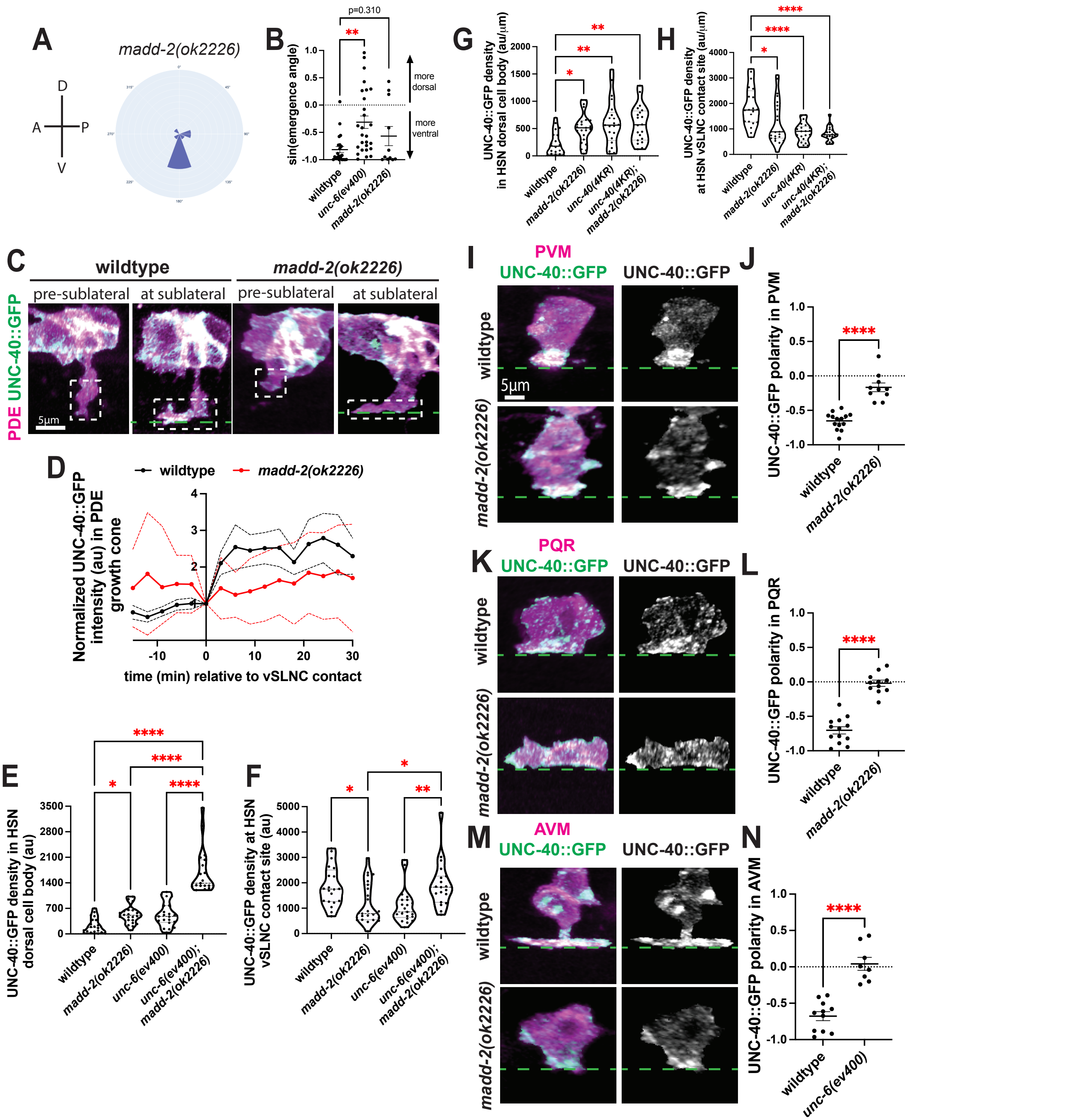


**Figure S2 UNC-40 polarization, but not axon emergence, is dependent on MADD-2, related to Figure 4.** A. Rose plot histogram of PDE axon emergence in *madd-2(ok2226*) animals. B. Graph comparing the sine function of axon emergence angle in wildtype, *unc-6(ev400)*, and *madd-2(ok2226)*animals. 1 indicates 90° dorsal axon emergence and -1 indicates 90° ventral axon emergence. C. Airyscan images of UNC-40::GFP expression PDE before and after vSLNC contact in wildtype and *madd-2(ok2226)* animals. Dashed box denotes growth cone. D. Normalized intensity traces of growth cone UNC-40::GFP during axon navigation normalized to sublateral contact (t=0). Black line denotes wildtype animals, and red line denotes *madd-2(ok2226)* animals. SEM is shown. E-F. Integrated UNC-40::GFP density along HSN dorsal cell body (E) and HSN contact site with vSLNC (F) in wildtype, *unc-6(ev400)*, *madd-2(ok2226),* and *unc-6(ev400); madd-2(ok2226)* animals. G-H. Integrated UNC-40::GFP density along HSN dorsal cell body (G) and HSN contact site with vSLNC (H) in wildtype, *madd-2(ok2226), unc-40(4KR)* and *unc-40(4KR); madd-2(ok2226)* animals. I,K,M. Confocal images of UNC-40::GFP distribution in PVM (I), PQR (K), and AVM (M) in wildtype and *madd-2(ok2226)* animals. J,L,N. Graphs of UNC-40::GFP polarity PVM (J), PQR (L), and AVM (N) in wildtype and *madd-2(ok2226)* animals. SEM is shown Scale bar is 5 µm in C,I,K,M. Ordinary one-way ANOVA with multiple comparisons is used in B,E-H. Unpaired t-test is used in J,L,N.

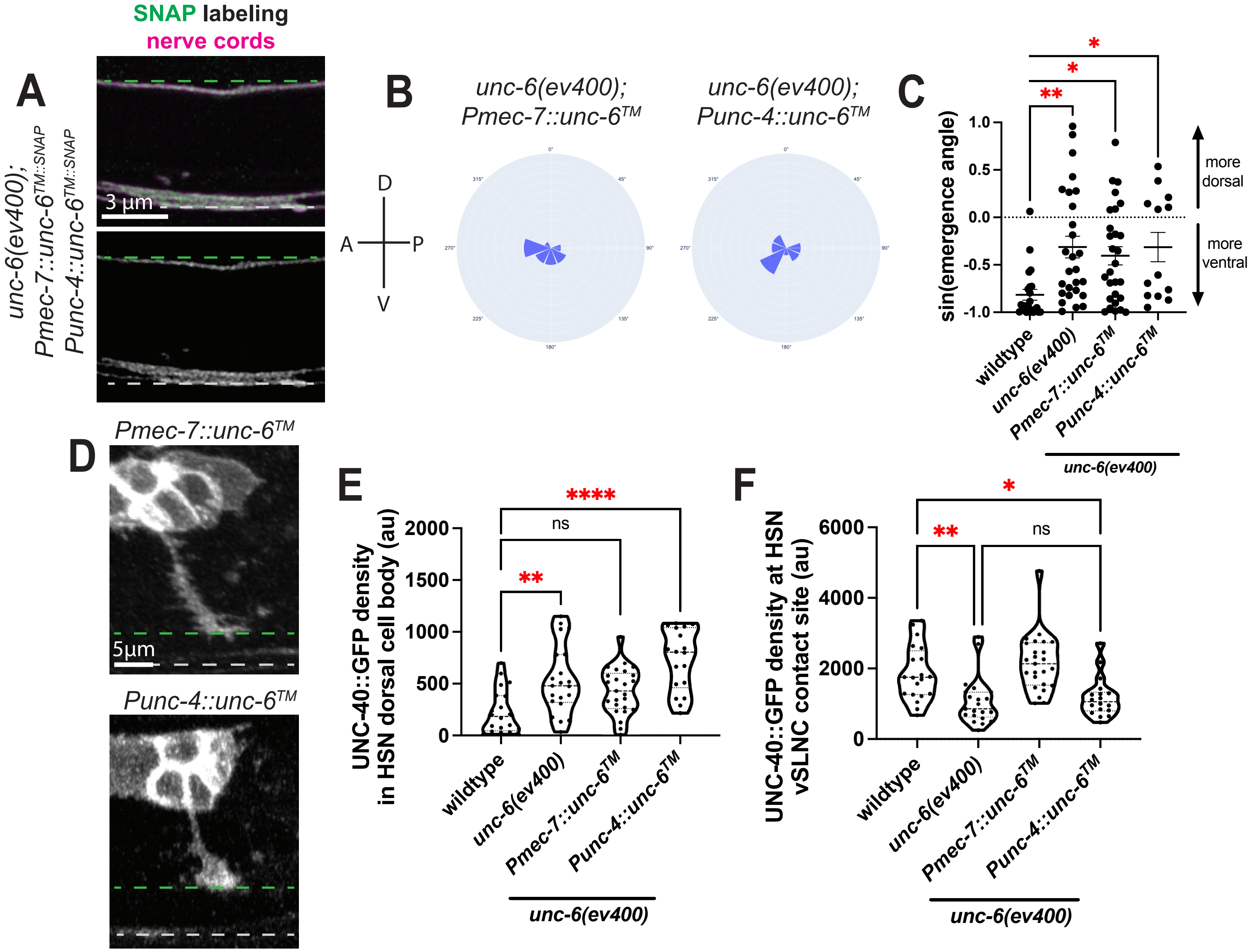


**Figure S3 Membrane tethered UNC-6 is not sufficient to explain axon emergence, related to Figure 5.** A. SNAP labeling of *unc-6(ev400); Pmec-7::unc-6$^{TM}$; Punc-4::unc-6$^{TM}$* animals. B. Rose plot histograms of PDE axon emergence in *unc-6(ev400); Pmec-7::unc-6$^{TM}$* and *unc-6(ev400); Punc-4::unc-6$^{TM}$.* C. Graph comparing the sine function of axon emergence angle in wildtype, *unc-6(ev400)*, *unc-6(ev400); Pmec-7::unc-6$^{TM}$* and *unc-6(ev400); Punc-4::unc-6$^{TM}$* animals. 1 indicates 90° dorsal axon emergence and -1 indicates 90° ventral axon emergence. D. Confocal images of *Pmec-7::unc-6$^{TM}$* and *Punc-4::unc-6$^{TM}$* animals showing normal PDE axon interactions with the vSLNC. E-F. Integrated UNC-40::GFP density along HSN dorsal cell body (E) and HSN contact site with vSLNC (F) in wildtype, *unc-6(ev400)*, *unc-6(ev400); Pmec-7::unc-6$^{TM}$* and *unc-6(ev400); Punc-4::unc-6$^{TM}$* animals. Scale bar is 3 µm in A and 5 µm in D. Ordinary one-way ANOVA with multiple comparisons is used in C,E,F.

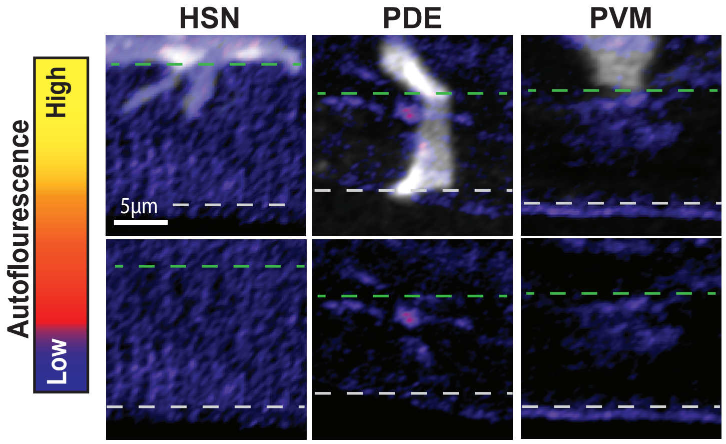


**Figure S4 Levels of autofluorescence in the region between the vSLNC and VNC are low, related to Figure 6.** Airyscan images of axon extension in HSN, PDE, and PVM in wildtype animals lacking endogenous UNC-6::mNG. Images were taken and processed with the same protocols as images in Figure 6A,B,D. Scale bar is 5 µm.
